# The anterior olfactory nucleus mediates curious exploration evoked by novel odors

**DOI:** 10.1101/2025.05.19.654779

**Authors:** R De Plus, M Broux, H Giaffar, E Schiltz, K Kondrakiewicz, C Aydin, S Haesler

**Affiliations:** Neuroelectronics Research Flanders (NERF), Leuven, Belgium; Department of Neurosciences, KU Leuven, Leuven, Belgium; Halıcıoğlu Data Science Institute, University of California, San Diego, La Jolla, USA; Department of Biomedical Engineering, Bahcesehir University, Istanbul, Turkey

## Abstract

The spontaneous exploratory reaction to novel stimuli reflects a fundamental form of curiosity, which is widely observed in the animal kingdom. How sensory systems mediate the recognition of novel stimuli to evoke exploration is not well understood. To address this question, we presented novel and familiar olfactory stimuli to head-restrained mice, while measuring novelty-evoked exploratory behaviors. In parallel, we recorded neural activity in primary olfactory cortical structures, the anterior olfactory nucleus (AON) or the anterior piriform cortex (aPCx). Novelty strongly modulated odor responses in the AON, but only weakly in the aPCx. Pharmacological and chemogenetic inactivation of the AON but not the aPCx disrupted exploratory responses. During long-term habituation over multiple days, sensory representations were drifting in the AON whereas they became stable within one day in the aPCx. Our findings suggest that AON and aPCx play distinct roles in novelty-evoked exploration. While the AON mediates the immediate reaction to novel stimuli, the aPCx exhibits stable stimulus representations, consistent with supporting odor memory.

## Introduction

In many species, novel or surprising stimuli evoke curious exploration (Modirshanechi et al., 2023). This spontaneous behavioral reaction involves short latency, orienting responses, including transient changes in skin conductance, respiration, pupil diameter and heart rate (Nieuwenhuis et al., 2011) and exploratory movements of sensory organs such as eyes or whiskers (Sokolov, 1963). Novel stimuli of different sensory modalities evoke broadly similar orienting responses (Scourse and Hinde, 1973; Inglis and Fibiger, 1995), suggesting the convergence of neural pathways for initiating orienting behaviors. The recognition of novelty, on the other hand, is performed by each sensory system separately. In the visual system, stimulus novelty enhances sensory responses as early as in the retina (Hosoya et al., 2005) but also at later processing stages in primary visual cortex (Garrett et al., 2020; Homann et al., 2022; Ito et al., 2024) through thalamo-cortical pathways (Furutachi et al., 2024). Damage to primary visual cortex disrupts orienting behaviors (Sprague, 1996). Stimulus novelty also increases sensory responses in the olfactory system, as early as in the olfactory bulb (OB) (Kato et al., 2012), but how novelty affects neural activity in primary olfactory cortical structures has not been investigated in awake animals. Moreover, it remains unknown if odor-related activity in primary olfactory cortices is critical for mediating orienting behaviors.

The most anterior cortical area to receive olfactory information from the OB is the anterior olfactory cortex (or nucleus, AON). The AON consists of two tangential layers; an outer tightly packed plexiform layer and an inner cell zone surrounding the anterior part of the anterior commissure (AC). Through the AC, the AON receives olfactory input from the contralateral OB, supporting its role in odor source localization (Rabell et al., 2017). AON feedback to the OB has further been shown to specifically control the gain of odorant responses while leaving odorant tuning unaffected (Quintela et al., 2020; Chae et al., 2022). The two layers of AON gradually merge with the three layers of the piriform cortex (PCx), traditionally referred to as primary olfactory cortex and the largest component of olfactory cortex. The PCx is characterized by a sparsely populated superficial layer, a dense input layer with glutamate-releasing principal neurons, and a deep layer with fewer principal neurons. Contrary to the AON, which receives input from both mitral and tufted cells in the OB, the PCx only receives input from mitral cells (Igarashi et al., 2012). Both the AON and PCx in turn provide extensive feedback to the OB through parallel processing loops, preferentially modulating tufted and mitral cells, respectively (Chae et al., 2022).

The PCx can be further divided into an anterior (aPCx) and a posterior (pPCx) section. While the aPCx is involved in odorant identification and discrimination (Wilson and Sullivan, 2011), the pPCx has been related to higher-order, associative functions, such as the creation of odor-place maps (Poo et al., 2022). Most studies on the neural representation of smell have focused on the aPCx. Odor representations in the aPCx are sparse and dispersed, integrating spatial and temporal inputs from the OB (Stettler and Axel, 2009; Bekkers and Suzuki, 2013). Odorant concentration in aPCx is reflected in the timing of spikes (Bolding and Franks, 2017), rather than spike rates (Roland et al., 2017; Bolding and Franks, 2018). Previous studies on the impact of experience on odorant representations in the olfactory cortices were conducted under anesthesia (Wilson, 1998, 2000, 2003), which has a major impact on olfactory processing (Wachowiak, 2011; Kato et al., 2012).

In this study, we recorded single-unit activity in the AON and aPCx while mice engaged in the spontaneous detection of novel odorants (Esquivelzeta Rabell et al., 2017; Mutlu et al., 2018). We used the stereotypical increase in respiratory frequency also referred to as exploratory sniffing (Wachowiak, 2011) as behavioral indicator of novelty detection. This allowed us to compare how novelty affects olfactory responses in the two main primary olfactory cortical structures. Moreover, we evaluated how responses evolved over multiple trials while behavioral responses habituated. We found that neuronal responses to familiar odorants were similar between AON and aPCx, but novel odorants differentially activated the AON and aPCx. Responses to novel odorants were larger in AON than aPCx. Moreover, responses in the AON decreased with recurrent stimulus exposure, in parallel to the behavioral habituation dynamics. In the aPCx, on the other hand, responses remained more consistent across trials. Importantly, the observed differences in response modulation cannot be explained by changes in sniffing frequency or net population firing rate but are a consequence of stimulus novelty. Finally, we demonstrate that acute pharmacological inactivation of the AON disrupts orienting behavior, while we observed little changes in orienting after inactivation of the aPCx. Consistent with our electrophysiologal recording results, behavioral responses were thus more sensitive to perturbation of the AON than the aPCx. Our findings establish that the AON is in the critical path for evoking exploration in response to novel odorants, which constitutes a fundamental form of curiosity.

## Results

### Neuronal responses to familiar odorants are comparable between AON and aPCx

To characterize olfactory responses in AON and aPCx, we performed high-density electrophysiological recordings in awake head-restrained C57Bl/6j mice, stimulated with odorants for 2 seconds. Before we started the recording sessions, animals were habituated to the behavioral setup and were familiarized with four different odorants over a period of 4 days (Fig 1a and Supplementary Fig. 1). All trials were aligned to the first inhalation after odorant onset. Odor-responsive units were then identified by statistic comparison (p<0.05, rank-sum test) of activity in a 400ms long time window after the first inhalation with baseline activity in a 400ms long time window 2s prior to odorant stimulation.

**Fig. 1.**
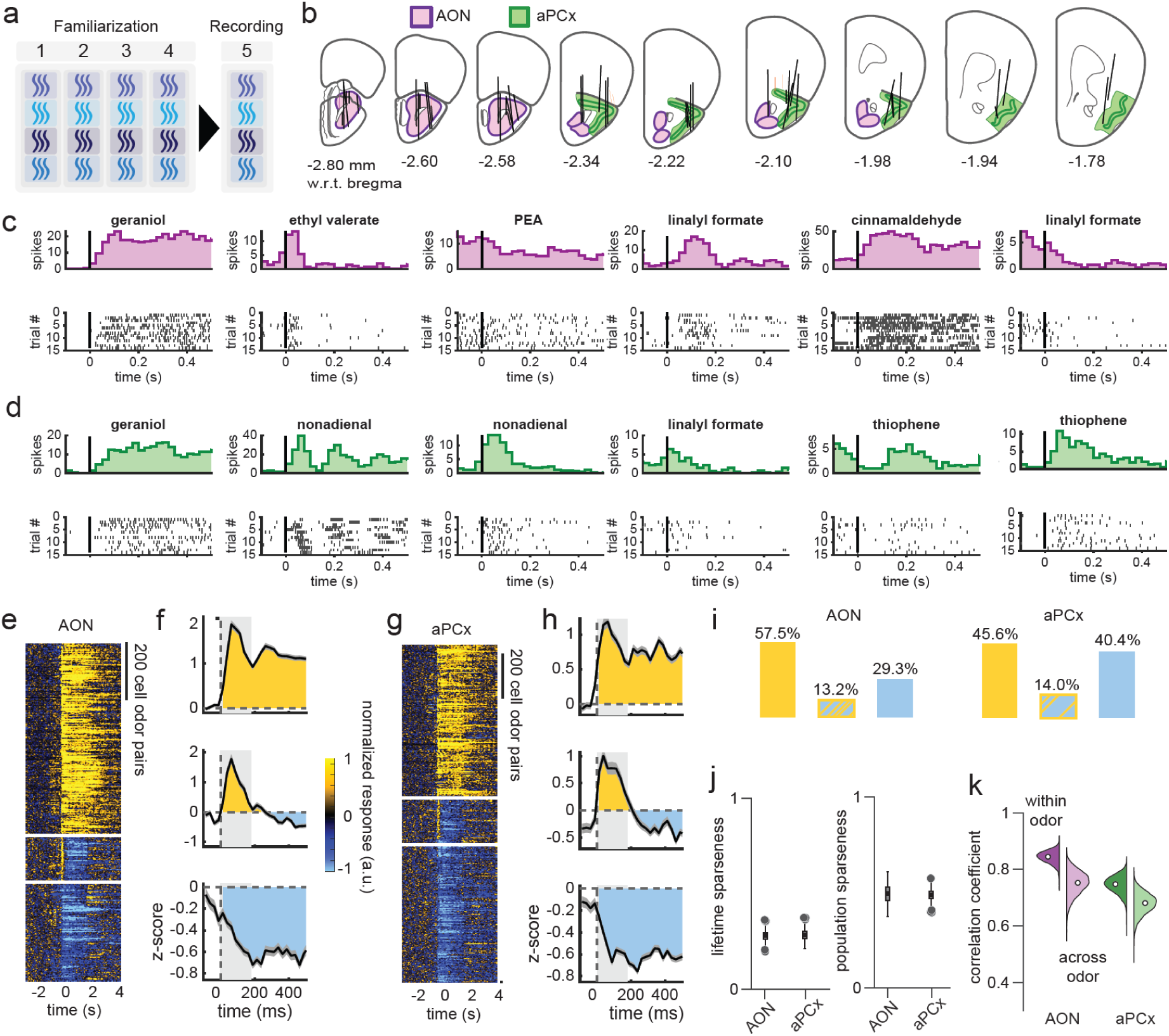
Neuronal responses to familiar odorants are comparable between AON and aPCx. **a** Experimental paradigm to record neuronal responses to familiar odors. Numbers indicate days of odor exposure. **b** Anatomical reconstruction of probe insertion trajectories (AON purple, aPCx green). **c** Spike raster plots (top) and peri-stimulus time histograms (bottom) of six example neurons in the AON, each responding to the indicated odorant (PEA: phenylethyl-alcohol). **d** Same as (d) recorded from aPCx. **e** Min-max normalized responses of AON cell odor pairs grouped by response type. **f** Mean response ± SEM of each group in (e). Positive and negative responses are labelled in yellow and blue, respectively. **g** Same as (e) for cell odor pairs from aPCx. **h** Same as (f) for aPCx. **i** Proportion of functional cell types (yellow: positive and sustained responses; yellow and dashed blue: transient increase and sustained decrease; blue: negative sustained responses) in AON (left) and aPCx (right). **j** Lifetime (left) and population (right) sparseness in AON and aPCx neurons. Grey dots represent outliers of the distribution (Kolmogorov–Smirnov test, ns: not significant, *p>*0.0.5). **k** Bootstrapped distributions of trial-to-trial correlations within and across scents in AON (left) and aPCx (right) were generated by selecting cells with replacement and calculating the mean correlation within and across odors 1000 times.

In total, we recorded 485/956 (responsive/total) well-isolated units from 19 separate recordings in 18 mice (Fig. 1b, Table 1). All trials were aligned to the first inhalation after odorant onset. Of all well-isolated neurons, 248 neurons in AON and 237 neurons in aPCx responded significantly to at least one odor. Odor-responsive cells in aPCx had relatively low baseline firing rates (mean: 6.6 spikes/s, st. dev.: 7.1 spikes/s, range: 0.8 and 25.5 spikes/s; 2.5% and 97.5%) and log-normal distribution of responses, as described previously (Bolding and Franks, 2017). In contrast, AON neurons showed higher firing rates (mean: 9.2 spikes/s, st. dev.: 10.2 spikes/s, range: 1.1 and 35.7 spikes/s, 2.5% and 97.5%, AON and aPCx, *p<*0.05, t-test on log transformed data), more similar to the olfactory bulb (Poo and Isaacson, 2009).

**Table 1:**
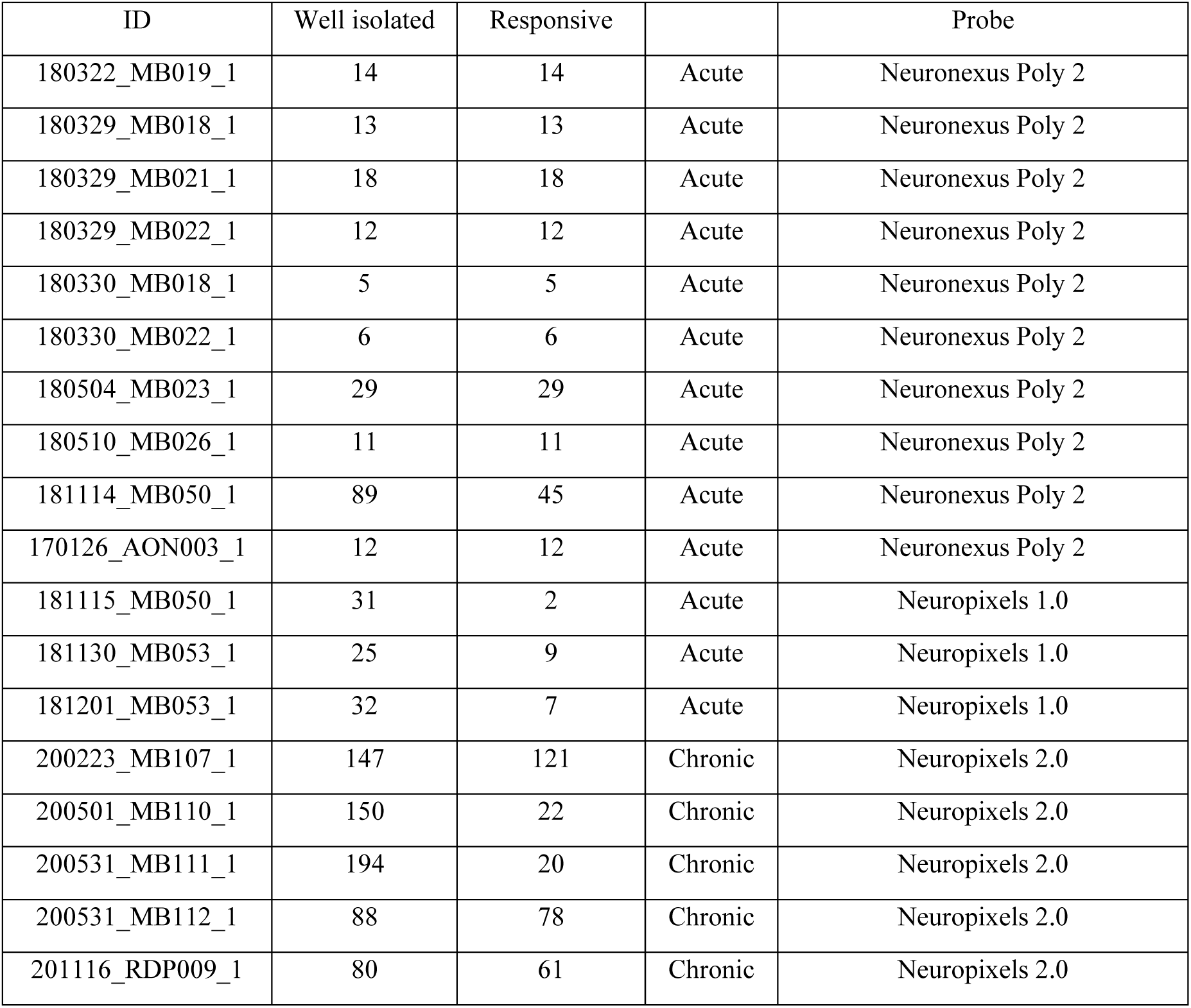
List of experimental animals in which electrophysiological recordings were performed.

| ID | Well isolated | Responsive |  | Probe |
| --- | --- | --- | --- | --- |
| 180322_MB019_1 | 14 | 14 | Acute | Neuronexus Poly 2 |
| 180329_MB018_1 | 13 | 13 | Acute | Neuronexus Poly 2 |
| 180329_MB021_1 | 18 | 18 | Acute | Neuronexus Poly 2 |
| 180329_MB022_1 | 12 | 12 | Acute | Neuronexus Poly 2 |
| 180330_MB018_1 | 5 | 5 | Acute | Neuronexus Poly 2 |
| 180330_MB022_1 | 6 | 6 | Acute | Neuronexus Poly 2 |
| 180504_MB023_1 | 29 | 29 | Acute | Neuronexus Poly 2 |
| 180510_MB026_1 | 11 | 11 | Acute | Neuronexus Poly 2 |
| 181114_MB050_1 | 89 | 45 | Acute | Neuronexus Poly 2 |
| 170126_AON003_1 | 12 | 12 | Acute | Neuronexus Poly 2 |
| 181115_MB050_1 | 31 | 2 | Acute | Neuropixels 1.0 |
| 181130_MB053_1 | 25 | 9 | Acute | Neuropixels 1.0 |
| 181201_MB053_1 | 32 | 7 | Acute | Neuropixels 1.0 |
| 200223_MB107_1 | 147 | 121 | Chronic | Neuropixels 2.0 |
| 200501_MB110_1 | 150 | 22 | Chronic | Neuropixels 2.0 |
| 200531_MB111_1 | 194 | 20 | Chronic | Neuropixels 2.0 |
| 200531_MB112_1 | 88 | 78 | Chronic | Neuropixels 2.0 |
| 201116_RDP009_1 | 80 | 61 | Chronic | Neuropixels 2.0 |

Olfactory cortex neurons showed diverse response properties (Fig. 1c-d). Some neurons in AON and aPCx increased their firing rates to odor stimulation, some decreased it, consistent with previous observations in the aPCx (Zhan and Luo, 2010; Bolding and Franks, 2017). Moreover, we observed neurons with a short-latency, transient firing rate increase which was followed by sustained inhibition (Fig. 1e-i). Neurons with similar dynamics have previously been documented in the locust olfactory system (Mazor and Laurent, 2005). To compare response types between AON and aPCx more systematically, we grouped cortical neurons from both regions according to the observed temporal response profiles using k-means clustering. Both cortical regions showed all three functional response types with varying distributions in each area (Fig. 1e-i).

Next, we analysed the receptive fields of odor-responsive neurons. To quantify the stimulus selectivity of individual neurons and neuronal populations in AON and aPCx, we calculated lifetime sparseness and population sparseness (Treves and Rolls, 1991), respectively. We found that neither measure was statistically distinguishable between AON and aPCx (Fig. 1j). To further characterize odorant tuning of neurons in AON and aPCx, we computed the similarity of odor responses within and across odors based on population firing rates for each odor, as previously described (Bolding et al., 2020). Overall, responses in both regions were correlated, showing a slightly higher but not significant response correlation for the AON population compared to the aPCx population (Fig. 1k). There was no inter-odor response separation in aPCx and AON, indicating that they were broadly tuned and primarily responding to multiple odors (Fig. 1k). We conclude that AON and aPCx neurons thus had broadly similar receptive fields.

### Novel stimuli differentially modulate odor responses in AON and aPCx

To investigate the impact of novelty on odorant representations in primary olfactory cortical structures in awake animals, we introduced 8 novel odorants randomly interleaved with 4 familiar odorants (Fig. 2a). As shown previously (Wachowiak, 2011), mice engaged in exploratory sniffing in response to novel odors, while familiar odors evoked only small changes in respiration (Fig. 2b and Supplementary Fig. 2a). Exploratory sniffing was observed across a wide range of structurally dissimilar molecules, demonstrating that behavioral responses to novel odorants were largely invariant to the chemical nature of the stimulus (Fig. 2e). Recurrent stimulus exposure resulted in the progressive reduction of sniffing responses, indicating behavioral short-term habituation (Fig. 2c). Since breathing can modulate neural firing, we focused all further analysis on a time window of 400ms from the first inhalation onset after odor onset during which respiration rate was indistinguishable between novel and familiar odors (Fig. 2d).

**Fig. 2.**
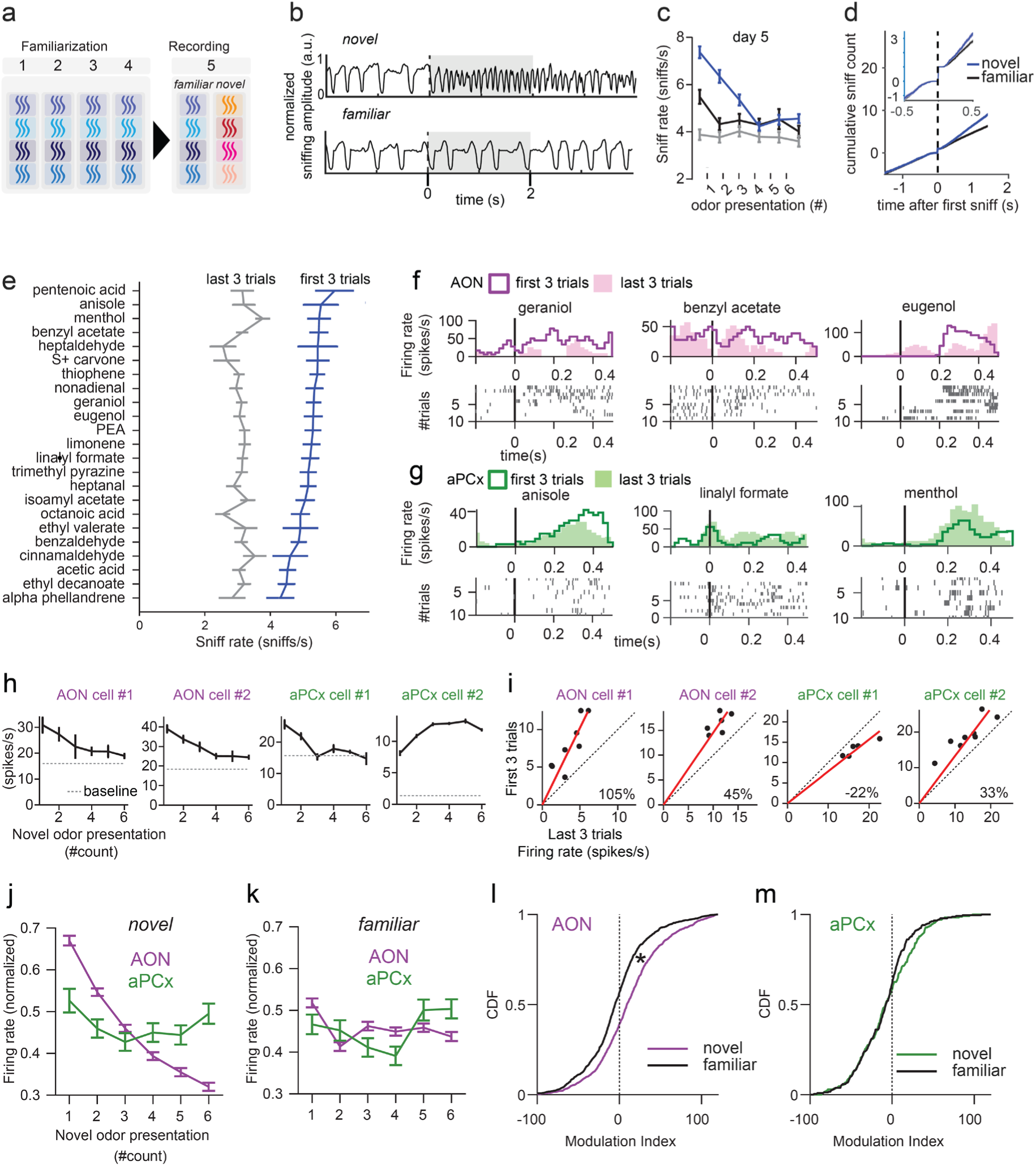
Novel stimuli differentially modulate odor responses in AON and aPCx. **a** Experimental paradigm to record neuronal responses to novel and familiar odors. Numbers indicate days of odor exposure. **b** Example sniffing traces extracted during exposure (grey shading) to a novel (top) and familiar (bottom) odorant. **c** Mean sniffing rate before odor stimulation (gray) and in response to familiar (black) and novel (blue) odors (mean+-SEM; n=5 mice). **d** Cumulative sniff counts during presentation of novel (blue) and familiar (black) odorants (mean ± SEM). Dashed line represents 400ms after the first sniff. Inlet represents the same data with the time interval of 0.5 before and after first sniff. **e** Mice (n=22) show similar exploratory sniffing behavior in response to a broad range of structurally dissimilar chemicals. Mean sniffing rate was calculated over the first three presentations to each odor. Blue; novel, gray; pre-odor/blank. (mean ± SEM). **f** Peri-stimulus spike rasters (bottom) and histograms (top) of three example AON neurons after introduction of a novel odor in trial 1 and during subsequent trials. The vertical black line indicates the first inhalation after odor onset. **g** Same as (f) for neurons from aPCx. **h** Habituation profile of 2 example neurons from AON (purple) and aPCx (green) neurons to 10 presentations of a novel odor. **i** Modulation index computed for all novel stimuli in example neurons from AON and aPCx. Each dot represents the mean firing rate of a neuron to an odor in the first three vs. last three trials. The red line indicates the constrained fit of the dots represented in the panels. Percentage (%) indicates an increase or decrease in firing rate between the first and the last three trials. **j** Population habituation profile for AON (purple, n=248 cells) and aPCx (green, n=237 cells) to novel odors. Single neuron habituation profiles to novel odor set were min-max normalized and averaged. **k** Same as (j) for familiar odors. **l** Cumulative modulation indices computed for novel (purple) and familiar (black) stimuli of AON neurons (Kolmogorov–Smirnov test \**p<*0.05). **m** Same as (l) for the population of aPCx neurons (green).

Recurrent odorant presentation modulated the magnitude of neuronal odorant responses in the AON and aPCx. In both structures, we found examples of neurons that increased or decreased their activity; some neurons remained unchanged compared to the first presentation of the respective stimulus (Fig. 2f-i). We calculated the population firing rate for both regions by averaging the responses of neurons to novel or familiar stimuli (Fig. 2j-k). Since the wide dynamic range of different neuronal responses might introduce biases in the calculation of population firing rates, we normalized population firing rates using min-max feature scaling to bring all the values into the range of [0,1]. We found a robust monotonic decrease of responses to recurring novel odorants in AON, which we did not observe in the aPCx (Fig. 2j). Remarkably, responses in AON were still changing when sniffing behavior was asymptotic (Fig. 2c). For familiar odors, the population firing rate was stable across multiple presentations in both AON and aPCx (Fig. 2k). Repeating the analysis of population means without normalization yielded the same result (Supplementary Fig. 2b, c). Importantly, the observed differences between AON and aPCx could not be accounted for by different stimulus selectivity, as both population sparseness and lifetime sparseness were similar between regions (Supplementary Fig. 2d).

To quantify the effect of novelty on individual neurons, we calculated a modulation index for each neuron, which was based on the comparison between the mean firing rate of the first three and the last three presentations of the same odor (Aydın et al., 2018). Importantly, this modulation index is not sensitive to possible differences in stimulus selectivity between presentations or neurons. When we examined the cumulative distribution function of the modulation index over all neurons, separately for novel and familiar stimuli (Fig. 2l-m; Supplementary Fig. 2a), we found that novel odors strongly modulated neuronal responses compared to familiar odors in the AON. However, in the aPCx, the distribution of modulation indices was similar between novel and familiar conditions (Kolmogorov-Smirnov test; *p<*0.05, AON novel vs familiar; *p>*0.05, aPCx, novel vs familiar). These results demonstrate that novelty differentially affected neural responses in the AON and aPCx.

### The effect of novelty on odor responses cannot be explained by sniffing frequency and net population firing rate

The behavioral reaction to novel stimuli was no longer detectable after three presentations. Thus, we predicted that the impact of novelty on neural representations was similarly transient and that neural representations would eventually converge to a stable representation during short-term habituation. To test this hypothesis, we measured population distances in neural activity space separately for AON and aPCx, using all trials that used the same odorants (Bolding et al., 2020).

We found that activity in both regions eventually stabilized with a recurring presentation of the same novel stimulus. However, the dynamic of this process was different between the two regions. In the AON, the distance to stable trials was larger and it took more trials to stabilize than in the aPCx (Fig. 3a, d; Supplementary Fig. 3a, e). This effect was not accounted for by an unspecific increase of firing rates due to general arousal, as the mean population firing rate averaged over all stimuli was relatively stable and similar between AON and aPCx (Fig. 3b, e; Supplementary Fig. 3b, f). Similarly, average sniffing rates during our analysis window (Fig. 3c, f; Supplementary Fig. 3c, g) were stable and comparable between the recordings in the two olfactory cortical structures. A multiple linear regression analysis using trial number, mean population firing rate and sniffing rate as independent variables confirmed that trial number had the strongest impact on neural activity (Fig. 3g-h; Supplementary Fig. 3d, h). However, in the AON (Fig. 3g; Supplementary Fig. 3d), the effect was substantially larger than in the aPCx (Fig. 3h Supplementary Fig. 3h).

**Fig. 3.**
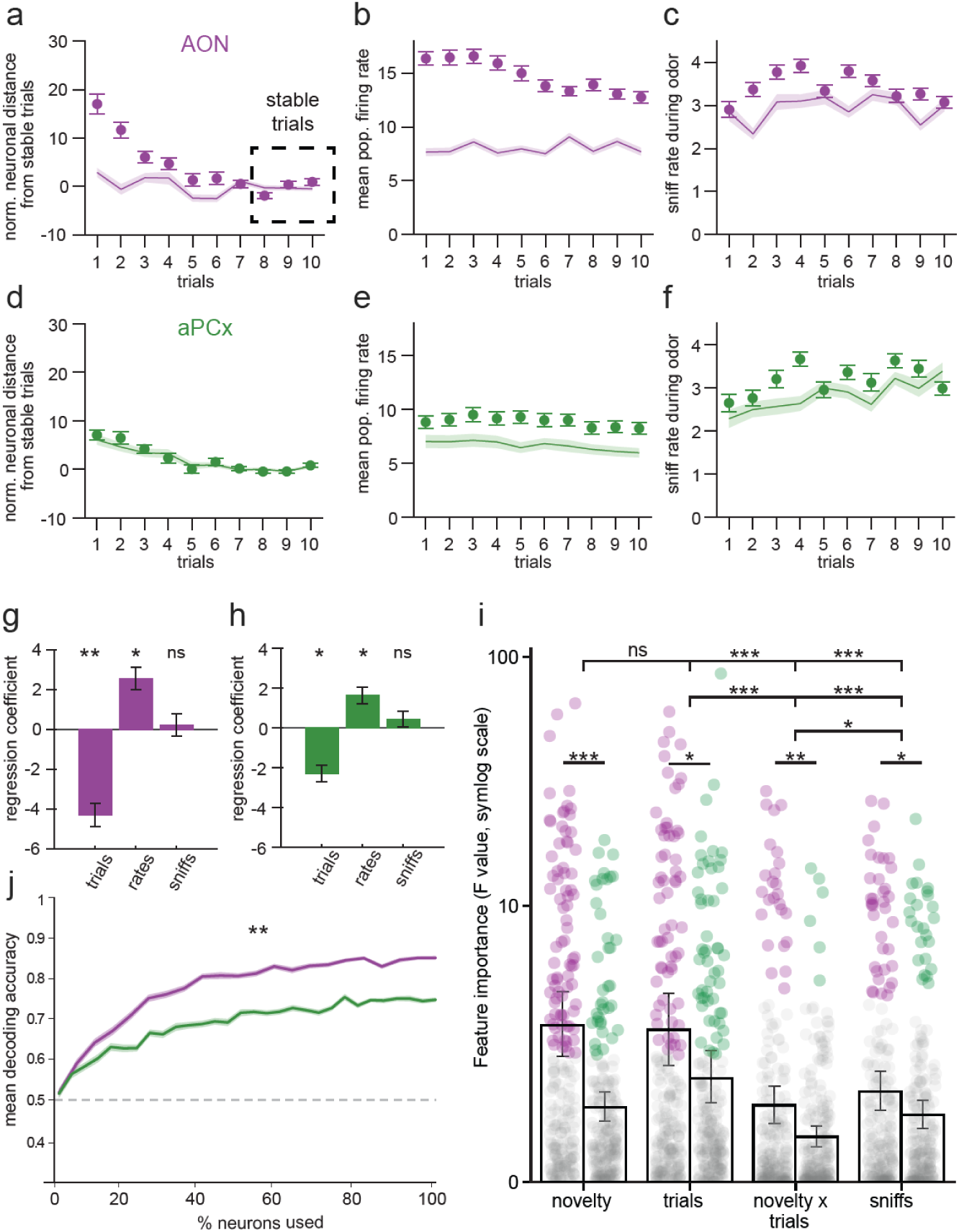
The effect of novelty on odor responses cannot be explained by sniffing frequency and net population firing rate. **a** Average Euclidean distance between novelty trial population vectors and stable trials in AON (n=258 cell odor pairs; shading indicates ± SEM). Baseline activity prior to odor presentation is represented as continuous line. **b** Average population firing rates to novel odors in AON (shading indicates ± SEM). Baseline activity prior to odor presentation is represented as continuous line. **c** Average AON sniff rates to novel odors (shading indicates ± SEM). Baseline respiration rates prior to odor presentation are represented as continuous line. **d-f** Same as (a-c) for aPCx (n=195 cell odor pairs). **g** Multiple linear regression coefficients for effects of trial number (\*\**p*<0.01), population firing rate (\**p*<0.05), sniff rate (*ns*: not significant) on population distance to stable in AON (mean ± SEM).**h** same as in (g) for aPCx (trial number and population firing rate \**p*<0.05, sniff rate *ns:* not significant). **i** Ordinary least squares linear regression with novelty, trial number, sniffing rate and an interaction term as predictors and mean firing rate in each trial as dependent variable in the full model. Stepwise exclusion of predictors (novelty, trials, novelty x trials, sniffs) revealed significant contributions of individual predictors (mixed ANOVA, F(3, 1320) = [19.85], p < 0.001; pairwise post-hoc comparisons t-tests followed by FDR correction for multiple comparisons \*\*\**p<*0.001, \*\**p<*0.01, \**p<*0.05) and between the two brain regions (mixed ANOVA F(1,440) = [24.14]). **j** Accuracy of the SVM in decoding novelty in AON (purple) or aPCX (green) as a function of the proportion of neurons used (Welch T-test, \*\**p*<0.01). The dashed line indicates chance level.

Complementary to the population analysis, we performed an ordinary least squares linear regression to quantify how much the response of each neuron was influenced by the experimental variables. The full model included odor novelty, trial number, sniffing rate and an interaction term as predictors and mean firing rate in each trial as dependent variable. To disentangle the individual contribution of these factors to neural firing dynamics, we stepwise excluded predictors and compared the resulting reduced models to the full model. This analysis revealed significant differences between the contribution of individual predictors and how they influenced the two brain regions (Fig. 3i). In particular, we found that novelty and trial number explained more variance than sniffing. Moreover, consistent with the population analysis, the effect of novelty was larger in the AON than in the aPCx.

Finally, we asked how well stimulus novelty was represented in the AON and aPCx using a linear Support Vector Machine (SVM). The accuracy of decoding novelty was above chance level in both AON and aPCx, but significantly higher in AON than in aPCx (Fig. 3j). These findings support the conclusion that novelty has a comparatively larger impact on odor responses in the AON than in aPCx.

### Odor responses in AON and aPCx follow different dynamics during long-term habituation

When we familiarized animals with stimuli over multiple days (Fig. 4a), we observed that sniffing responses to the first stimulus presentation gradually decreased. This behavioral long-term habituation was superimposed with daily short-term habituation, indicating two non-associative learning processes occurred in parallel.

**Fig. 4.**
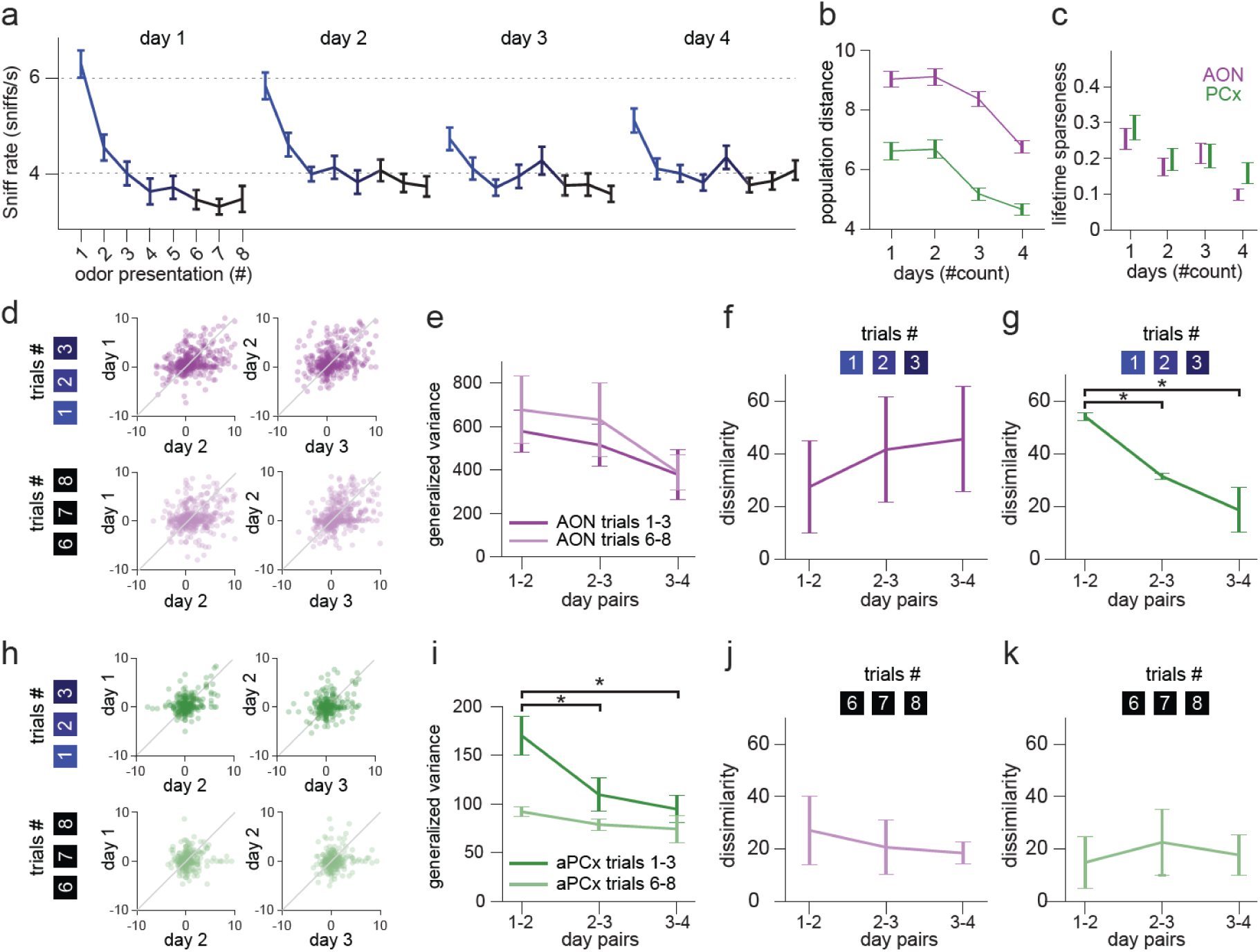
Odor responses in AON and aPCx follow different dynamics during long-term habituation. **a** Mean sniffing rate in response to odorants presented on 4 consecutive days. (mean+-SEM; n=7 mice). **b** Average Euclidean distance between novelty trial population vectors on different days and stable trials on day 4 in AON (purple, n=464 chain cell odor pairs; shading indicates ± SEM) and aPCx (green, n=404 chain cell odor pairs; mean ± SEM). **c** Lifetime sparseness of chain neurons in AON (purple) and aPCx (green), with error bars denoting mean ± SEM. **d** The mean spike counts of AON chain cell-odor pairs in the first three trials (top, dark purple) and trials 6-8 (bottom, light purple) are compared between day 1/day 2 and day 2/day 3. **e** Comparison of generalized variance, a measure of the variability of the point clouds in d between days (day pairs; dark purple: first three trials; light purple: trials 6-8) in in AON. **f** We used a CKA-based dissimilarity index to compare the resemblance of neural representations in the first three trials between different days in the AON (dark purple, n=3 mice). Low dissimilarity values indicate two representations resemble each other, whereas large values indicate their dissimilarity. The dissimilarity index was computed over chain neurons for each mouse. **g** Dissimilarity index for the first three trials between different days in aPCx (dark green, n=4 mice, ANOVA \**p<*0.05). **h** The mean spike counts of aPCx chain cell-odor pairs in the first three trials (top, dark green) and trials 6-8 (bottom, light green) were compared between day 1/day 2 and day 2/day 3. **i** Comparison of generalized variance between days (dark purple: first three trials; light purple: trials 6-8, ANOVA \**p<*0.05) in aPCx. **j** CKA-based dissimilarity in trials 6-8 between different days in AON (left, light purple) **k** same as j but for aPCx (right, light green).

To investigate how responses to the same odor evolved during long-term habituation, we tracked the activity of individual neurons over days. Using an algorithm based on Earth Mover Distance (Yuan et al., 2024), we successfully identified a total of 217 (AON: n=116, aPCx: n=101) “*chain neurons*” which could be consistently recognized on consecutive days (see Methods). We first applied the analysis of population firing rates (Fig. 3) to the chain neurons, normalizing responses of each day to the stable trials (last 3 trials) of the fourth day (day 4) of familiarization. We found that population firing rates in AON and aPCx decreased similarly to the observed behavioral habituation (Fig. 4b). Odor selectivity, as measured by lifetime sparseness, decreased slightly over multiple days, indicating that neurons in AON and aPCx maintained their broad odor tuning during long-term habituation (Fig. 4c).

Next, we compared the specific odor response of each neuron between all days. We separately analyzed spike counts of all cell-odor pairs in the first three trials (trials # 1-3, Fig. 4d, h, top) and the last three trials (trials # 6-8, Fig. 4d, h, bottom) to capture the effect of long-term habituation before and after behavioral responses had diminished because of short-term habituation (Fig. 4a). We found that the responses of neurons differed between days in both AON and aPCx as indicated by the spread of spike counts around the unity line (Fig. 4d, h). To quantify the magnitude of this variability, we calculated the generalized variance (GV) of each point cloud (Fig. 4e, i). We found that cell-odor pairs in AON exhibited large day-to-day variability, both in the first three and the last three trials (Fig. 4e). In aPCx on the other hand, variability of the first three trials consistently decreased (Fig. 4i, dark green), whereas it was stable from day 1 for the last three trials (Fig. 4i, light green). These findings suggest odor responses of AON neurons maintain a large variance across days, which only slowly decreases with long-term habituation. On the contrary, in aPCx the variance odor responses rapidly stabilized with long-term habituation (Fig. 4i).

While GV provides a suitable measure of the changes of neural responses over time, it is sensitive to differences in overall firing rates between regions, which prohibits the comparison of AON and aPCx. To address this limitation, we used a dissimilarity index based on Centered Kernel Alignment (CKA) to quantify the similarity between odor representations on different days and between different regions. The CKA measure focuses on the geometry of the neural representation, independent of its magnitude, providing a more robust measure for cross-regional comparisons. Consistent with our earlier observations, we found odor representations in AON consistently drifted from the first day to the third day, irrespective of whether the first three (Fig. 4f) or the last three trials (Fig. 4j) were analyzed. Remarkably, the odor representations in aPCx trials stabilized within one day (Fig.4g) and remained stable thereafter (Fig. 4k).

### The AON is necessary for eliciting exploratory behaviors induced by novel olfactory stimuli

Our electrophysiological recordings revealed that odor responses in the AON were strongly modulated by novelty, while they were more invariant to the effect of experience in the aPCx. Therefore, we hypothesized that the AON would be critical for evoking olfactory exploratory behaviors in response to novel odorants. To test this hypothesis, we used a microcatheter (Marcigaglia et al., 2024) chronically implanted into the AON to perform injections with either the GABA_A_ receptor agonist muscimol or artificial cerebrospinal fluid (aCSF) in behaving animals (Fig. 5a). We found that the injection of muscimol disrupted behavioral responses to novel stimuli, while the injection of aCSF had no effect (Fig. 5a, b). Importantly, muscimol did not affect the respiration during presentation of familiar odorants (Fig. 5a, c) nor baseline respiration rates (paired t-test *p*=0.72, data not shown). To investigate whether the effect of AON inhibition was selective to respiratory novelty responses, we also examined pupil reactions as a second, non-respiratory measure of exploratory orienting. We found that muscimol injection into the AON reduced the pupillary reaction to novel stimuli (Supplementary Fig. 5c, d), but not to familiar stimuli (Supplementary Fig. 5c, e). When we performed muscimol injections into the aPCx, we found neither an effect on novelty-evoked exploratory sniffing (Fig. 5d, e) nor on the pupillary reaction (Supplementary Fig. 5f, g). Also, respiration and pupillary reaction during presentation of familiar odorants (Fig. 5d, f*; S*upplementary Fig. 5f, h) was affected. Importantly, both AON and the anterior section of the aPCx were injected with the same volume of muscimol at the same rate, resulting in similar volumes of tissue coverage (Supplementary Fig. 5a, b).

**Fig. 5.**
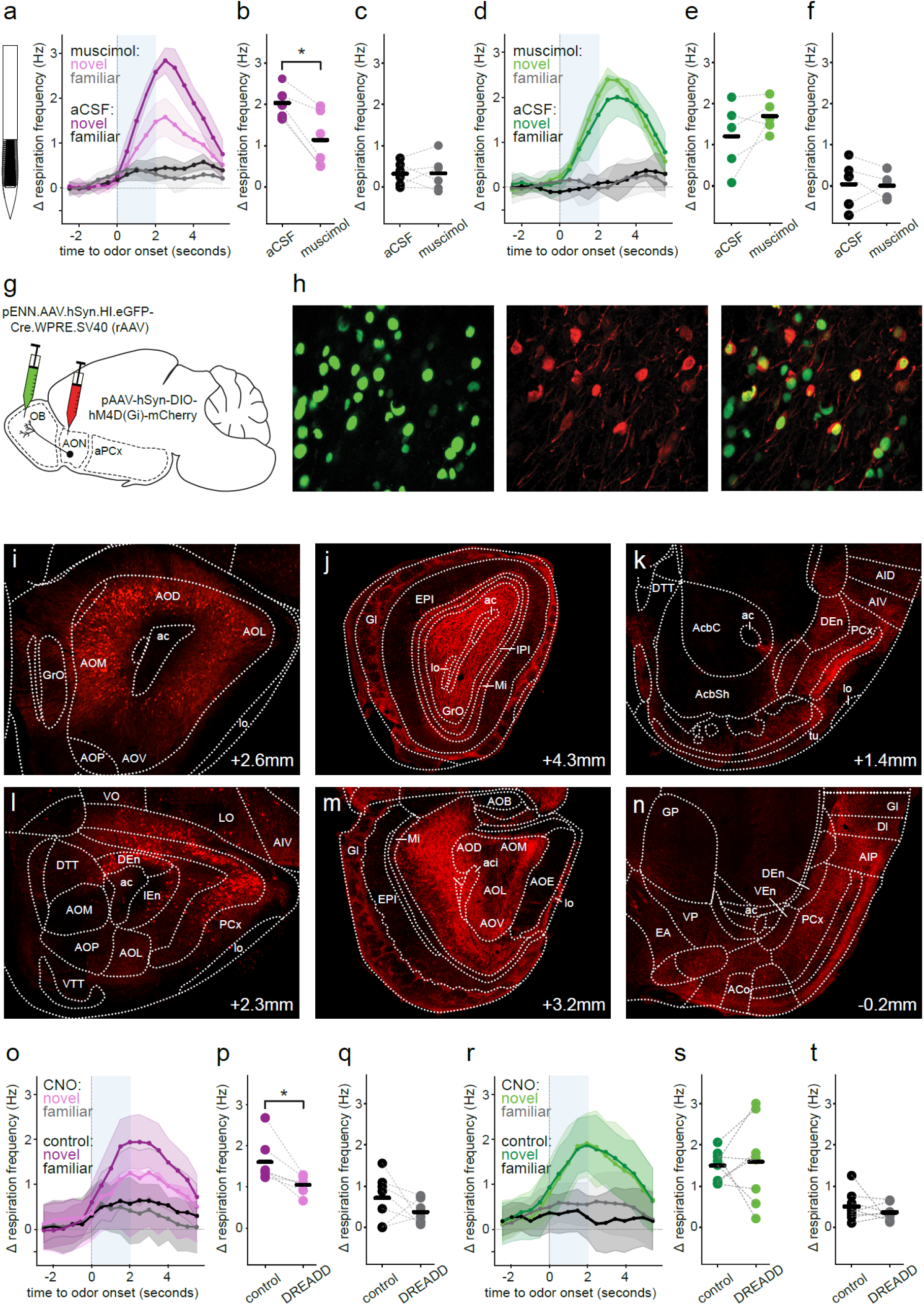
The AON is necessary for eliciting exploratory behaviors induced by novel olfactory stimuli. **a** Illustration of the microfabricated catheter used for convection-enhanced delivery of muscimol (left) and normalized change in breathing frequency during the first three trials (right, mean ± SEM) after AON injection with muscimol (novel trials: light purple, familiar trials: light grey) or aCSF (novel trials: dark purple, familiar trials: dark grey, n=5 mice). Blue shading represents the odor stimulation window. **b** Mean response (normalized change in breathing frequency during the time interval 0.5-3s in the first three trials) to novel stimuli after injection of muscimol or aCSF in the AON (paired t-test, \**p*=0.02). **c** Same as (b) for first three familiar trials (paired t-test, *p*=0.89). **d** Normalized change in breathing frequency during the first three trials (mean ± SEM) after aPCx injection with muscimol (novel trials: light green, familiar trials: light grey) or aCSF (novel trials: dark green, familiar trials: dark grey, n=5 mice). **e** Mean response (normalized change in breathing frequency during the time interval 0.5-3s in the first three trials) to novel stimuli after injection of muscimol or aCSF in the aPCx (paired t-test, *p*=0.26). **f** Same as (e) for the first three familiar trials (paired t-test, *p*=0.81). **g** Overview of the viral strategy to target OB-projecting neurons in the AON for chemogenetic inhibition. Retrograde AAV expressing Cre recombinase and GFP (pENN.AAV.hSyn.HI.eGFPCre.WPRE.SV40) was injected into the OB and AAV expressing inhibitory DREADD and mCherry (pAAV-hSyn-DIOhM4D(Gi)-mCherry) in the AON. **h** Immunohistochemical staining for GFP (left), mCherry (middle) and overlay of both (right). **i-k** Immunohistochemical staining for mCherry in brain slices to visualize DREADD expression after injection of DREADD virus into the AON. Anterior-posterior coordinates are indicated in the lower right corner of each image. **i** Expression in cell bodies of neurons in the AON. **j** Expression in axons of the OB (feedback projections). **k** Expression in axons of aPCx (feedforward projections). **l-n** Visualization of DREADD expression after injection of DREADD virus into the aPCX. **l** Expression in cell bodies of neurons in the aPCx **m** Expression in axon in the OB (feedback projections) **n** Expression in axons in the pPCx (feedforward projections). **o** normalized change in breathing frequency during the first three trials (mean ± SEM) after injection of CNO (novel trials: light purple, familiar trials: light grey) or saline (novel trials: dark purple, familiar trials: dark grey) in animals (n=7 mice) with DREADD virus targeted to AON. **p** Mean response (normalized change in breathing frequency during the time interval 0.5-3s in the first three trials) to novel stimuli after injection of CNO or saline in animals with DREADD virus targeted to AON (paired t-test, \**p*=0.02). **q** Same as (p) for first three familiar trials (paired t-test, *p*=0.29). **r** normalized change in breathing frequency during the first three trials (mean ± SEM) after injection of CNO (novel trials: light green, familiar trials: light grey) or saline (novel trials: dark green, familiar trials: dark grey) in animals (n=7 mice) with DREADD virus targeted to aPCx. **s** Mean response (normalized change in breathing frequency during the time interval 0.5-3s in the first three trials) to novel stimuli after injection of CNO or saline in animals with DREADD virus targeted to aPCx (paired t-test, *p*=0.90). **q** Same as (p) for first three familiar trials (paired t-test, *p*=0.68). **Abbreviations:** anterior commissure (**ac**), accumbens nucleus, shell (**AcbSh**), accumbens nucleus, core (**AcbC**), anterior cortical amygdaloid area (**Aco**), dorsal agranular insular cortex (**AID**), posterior agranular insular cortex (**AIP**), ventral agranular insular cortex (**AIV**), accessory olfactory bulb (**AOB**), dorsal/lateral/medial/ventral/posterior anterior olfactory nucleus (**AOD/AOL/AOM/AOV/AOP**), anterior olfactory nucleus pars externa (**AOE**), dorsal endopiriform claustrum (**DEn**), dorsal tenia tecta (**DTT**), extended amygdala (**EA**). external plexiform layer of the olfactory bulb (**EPI**), glomerular layer of the olfactory bulb (**Gl**), globus pallidus (**GP**), granular insular cortex (**GI**), granule cell layer of olfactory bulb (**GrO**), intermediate endopiriform claustrum (**IEn**), internal plexiform layer of olfactory bulb (**IPI**), lateral olfactory tract (**lo**), lateral orbitofrontal cortex (**LO**),mitral cell layer of the olfactory bulb (**Mi**), piriform cortex (**PCx**), olfactory tubercle (**tu**), ventral endopiriform claustrum (**VEn**), ventral pallidum (**VP**), ventral orbitofrontal cortex (**VO**), ventral tenia tecta (**VTT**).

In complementary manipulation experiments, we used a chemogenetic approach to directly inactivate olfactory cortical pyramidal cells. Given the rapid and profound influence of cortical feedback on OB output (Markopoulos et al., 2012), we specifically hypothesized that neurons projecting to the OB might play a prominent role in shaping odor responses during short-term habituation. To target OB-projecting neurons, we used a viral strategy involving a combination of two AAV. We injected retrograde AAV expressing the Cre recombines tagged with a green fluorophore into the OB, and anterograde AAV expressing Cre-conditional DREADD receptors tagged with a red fluorophore in the AON. Immunohistochemical analysis confirmed that the expression of the DREADD receptor in the AON was confined to neurons expressing Cre-recombinase that project to the OB (Fig. 5g). Cells expressing the DREADD also expressed Cre-recombinase (94.4% ± 0.6%; mean ± SEM, n=250 cells, Fig. 5h). In addition to widespread labelling of cell bodies in the AON (Fig. 5i), we also observed strong fluorescent signals in AON axons projecting to the OB (Fig. 5j) and the aPCx (Fig. 5k), suggesting that a considerable fraction of pyramidal cells in the AON simultaneously delivers both feedback to the OB and feed-forward signals to the aPCx. Of note, this does not preclude the existence of a population of neurons which exclusively project to the OB.

Inactivation of this central population of AON pyramidal cells disrupted exploratory sniffing responses (Fig. 5o, p) and pupil dilatation to novel stimuli (Supplementary Fig. 5i), while neither baseline respiration rates (data not shown) nor respiratory (Fig. 5q) and pupillary responses to familiar odors (Supplementary Fig. 5j) were affected. We also ruled out that the chemogenetic ligand clozapine N-oxide (CNO) influenced behavior on its own. Mice not expressing the DREADD receptors injected with saline or CNO showed indistinguishable baseline breathing rates (Supplementary Fig. 5m) as well as similar behaviors during presentation of responses to novel and familiar odors (Supplementary Fig. 5n, o).

For comparison, we also targeted aPCx pyramidal cells with our viral strategy. We injected retrograde AAV expressing Cre recombines tagged with a green fluorophore into the OB, and anterograde AAV expressing Cre-conditional DREADD receptors tagged with a red fluorophore in the aPCx. This approach resulted in widespread labelling of cell bodies in the aPCx (Fig. 5l), and strong fluorescent signals in aPCx axons projecting to the OB and the AON (Fig. 5m) as well as posterior PCx (Fig. 5n). A considerable fraction of pyramidal cells in the aPCx thus delivers both feedback to the OB and the AON. Inactivation of this cell population did not alter sniffing (Fig. 5r, s) or pupillary responses (Supplementary Fig. 5k) to novel odors. Baseline respiration rates (data not shown) as well as respiration (Fig. 5r, t) and pupil diameter (Supplementary Fig. 5l) during the presentation of familiar odors were also unaffected by the manipulation. Importantly, the number of cells labelled by our viral strategy in the AON and aPCx, respectively, was similar (AON n=699 ± 51 cells, aPCx n=669 ± 23 cells; mean ± SEM, t-test not significant, see Methods). Collectively, our results reveal a functional difference between the AON and aPCx and establish that the AON is critical for mediating exploration in response to olfactory novelty.

## Discussion

In this study, we investigated how olfactory cortices AON and aPCx mediate the recognition of novel stimuli to evoke sensory exploration. To address this question, we performed large-scale electrophysiological recordings in head-restrained mice exposed to novel and familiar stimuli. Mice spontaneously reacted to novel but not familiar stimuli with exploratory sniffing behavior, which habituated rapidly with recurring presentation of a novel stimulus. While familiar odorants evoked largely similar responses in the AON and aPCx, novel stimuli differentially modulated neuronal responses in AON and aPCx. In the AON, novelty increased the magnitude of odor responses, while not affecting stimulus selectivity. On the other hand, responses in the aPCx were more invariant to the effect of experience. Importantly, the impact of novelty on neural representations could not be explained by a modulation of sniffing frequency or net population firing rate. When we tracked neuronal responses over multiple days during long-term habituation, we observed sensory representations were drifting in the AON whereas they became stable within one day in the aPCx. Finally, we evaluated the functional role of the olfactory cortex in novelty-evoked exploratory sniffing. Pharmacological inactivation of the AON but not the aPCx disrupted exploratory sniffing induced by novel smells. Chemogenetic inactivation of AON pyramidal cells projecting to OB similarly reduced exploratory sniffing. Both manipulations also affected pupillary novelty responses, suggesting that the AON is in the critical path for evoking olfactory novelty-evoked exploration in general.

Our findings uncover a differential contribution of AON and aPCx to processing novelty and evoking exploratory behaviors. How do the physiological and functional differences between AON and aPCx relate to their differential anatomical organization? Whereas both structures receive input from mitral cells in the OB, the AON also receives input from tufted cells (Igarashi et al., 2012). The effect of novelty on tufted cell firing has yet to be described, but mitral cells show much larger responses to novel than to familiar stimuli (Kato et al., 2012). This observation not only suggests the OB as the site of novelty recognition, but it might also largely explain the strong novelty-modulation of activity in the AON. Conversely, the relatively smaller impact of novelty on aPCx firing is inconsistent with novelty-modulated mitral cell inputs. The aPCx must have a mechanism to create a neural representation which is rather invariant to the effect of novelty. A possible circuit mechanism might involve the recurrent circuitry of the aPCx, which has been shown to recruit global feedback inhibition to normalize responses across different concentrations (Bolding and Franks, 2018; Bolding et al., 2020). Further experiments are needed to test the hypothesis that the aPCx creates an experience-invariant representation in a similar manner to creating a representation that is invariant to odor concentrations.

Both AON and aPCx provide extensive feedback to the OB through parallel processing loops, preferentially modulating tufted and mitral cells, respectively (Chae et al., 2022). AON feedback specifically modulates the gain of tufted cell responses without altering odor tuning (Chae et al., 2022). By boosting odor responses in the AON, novelty might therefore in turn enhance the gain of tufted cells in the OB and contribute to the plasticity underlying short-term habituation. Our finding that chemogenetic inactivation of the AON disrupts novelty-evoked orienting responses supports this hypothesis. However, our viral strategy to target OB feedback, did not disambiguate the contribution of OB feedback from a possible contribution of feedforward collaterals. To address this limitation, future research needs to determine whether all AON cells which provide feedback to the OB also have collateral projections to the aPC, or if a distinct population of AON pyramidal cells exists that exclusively project feedback to the OB.

Although we find no direct evidence for a major contribution of the aPCx to mediating novelty-evoked exploration, this does not preclude the possibility it plays a critical role. First, our experiments focused on the most anterior section of the aPCx. Moreover, due to a sparsening of the representations, novelty might specifically modulate a small, specialized population of neurons in the aPCx (Schiltz et al., 2024), which may be difficult to target by manipulation. On the other hand, it is possible that the path from novelty recognition to eliciting behavioral responses does not involve feedforward projections through the aPCx. Alternative routes could include direct projections from the OB or AON to the cortical amydala, olfactory tubercle, or mediodorsal thalamic nucleus (Brunjes et al., 2005; Courtiol and Wilson, 2015; Chen et al., 2022). It will be interesting to determine how these olfactory regions downstream of AON and aPCx are modulated by novelty.

To complement our analysis of odor representations during short-term habituation after novel stimuli were introduced, we also investigated odor representations during long-term habituation over multiple days. Consistent with earlier observations (Schoonover et al., 2021), the aPCx showed overall low variance of odor responses and odor representations between days (Fig. 4). The rapid stabilization of odor representations in aPCx after only one day of stimulus exposure indicates that the aPCx follows a one-shot-learning principle. On the contrary, the AON showed large drift and a slower stabilization of odor representations. Taken together, our results thus suggest a differential contribution of AON and aPCx to novelty-evoked exploration. Whereas the AON mediates the immediate reaction to novel or otherwise salient stimuli, the aPCx exhibits a stable odor representation, consistent with supporting long-term odor memory.

How do our findings relate to observations made in other sensory modalities? Olfaction is unique among the senses in that sensory information is directly routed from the primary sensory organ to cortical brain areas without first being passed to subcortical structures (Shepherd, 2003). In the auditory, visual and somatosensory systems, such subcortical structures as well as their cortical projection targets are critical for mediating rapid orienting reactions to novel, unexpected or otherwise salient stimuli (Sprague, 1996; Malhotra et al., 2008). Moreover, responses in primary sensory neocortices have been shown to be enhanced by novelty (Garrett et al., 2020; Hamm et al., 2021; Homann et al., 2022; Gong et al., 2024). In the olfactory system, we found that these properties observed in subcortical and primary sensory neocortical regions were similarly observed in the AON but were much less evident in the aPCx, although the aPCx is commonly referred to as primary olfactory cortex. In the neocortex, novelty can enhance sensory responses in a stimulus-specific manner through cortico-thalamic loops (Furutachi et al., 2024). A similar stimulus-selective gain-control mechanism might be implemented in feedforward-feedback loops between AON and the OB, which are organized in topographic manner (Ghosh et al., 2011; Miyamichi et al., 2011; Chae et al., 2022; Chen et al., 2022). Further research is needed to test the hypothesis that the evolutionary older olfactory system uses conserved principles of novelty processing which are implemented in an anatomically distinct manner in the other sensory modalities after more recent evolutionary innovations.

## Materials and methods

### Mice

All procedures were executed per the guidelines the Federation of European Laboratory Animal Science Associations set forth and received approval from the Animal Ethics Committee at KU Leuven (protocols P067/2017 and P073/2019). The experiments were carried out in a cohort of 18 male C57Bl/6j mice obtained from the KU Leuven animal facility, weighed between 22 and 30 g, and were 6 and 12 weeks old.

### Surgical procedures

All experimental animals were surgically implanted with a titanium head plate to permit head-restraining during behavioral experiments. Mice were anesthetized with an intraperitoneal injection of a mixture of ketamine (75mg/kg) and medetomidine (Domitor, 1mg/kg) and fixated on a feedback-controlled heating pad at 38°C in a stereotactic set-up (SR-6M-HT, Narishige). 50µl of Lidocaine (Xylocain) was injected subdermally for local anesthesia. A patch of skin on top of the skull was removed for acute electrophysiology surgeries so that lambda and bregma were visible. A dental drill (Athena Champion AC 5000) with a carbide drill head of 0.8mm diameter (FG1, NeoBurr) was used to roughen the skull, and a bone screw was inserted into the skull (M1, length: 2mm). The bone screw served as electrical ground for electrophysiological recordings. Finally, a head plate was fixated on the skull with dental cement (Super-Bond, [3-component: Quick Monomer, C&B Polymer L-type Radiopaque, Catalyst V], Sun Medical). A second layer of cement (Jet Denture Repair, Lang) was applied on top to increase strength. The following surgical steps were different for each experimental procedure described below.

### **i.)** Procedure for acute electrophysiological recordings

Above the target coordinates (AON: AP2.6 ML1.4; aPCx: AP2.0 ML1.3), a craniotomy was created and filled with artificial dura (Cambridge Neurotech) and closed off with silicone (Kwik-Sil, World Precision Instruments).

### **ii.)** Procedure for chronic probe implantation

For chronic Neuropixels 2.0 probe insertion surgeries, a 2mm long, 1.5mm wide craniotomy was made at the insertion target. A durotomy followed the craniotomy if penetration with dura was not possible. The Neuropixels 2.0 probe was dyed with DiI and the implant inserted as described previously (van Daal et al., 2021). The probe was fixed in place with an additional layer of cement (Super-Bond, see above), the reference screw was be connected to the implant reference, and a third layer of cement was added (Wave A2, light-curable, SDI; curated with radii-cal, SDI).

### **iii.)** Catheter implantation

For catheter implantations, the skin and connective tissue were removed over the target coordinates. The coordinates of the craniotomy were marked (AON: AP 2.7, ML 1.0; aPCx: AP 2.245, ML 1.75, Paxinos and Keith B. J. Franklin, 2007). Polyacrylamide (Vetbond) was used at the edges to limit skin irritation due to cement and the chance of infection. Using a dental drill, ridges were created in the skull to broaden the surface area for more adhesion when dental cement was applied. After performing a bilateral craniotomy, a custom microfabricated brain catheter (Marcigaglia et al., 2024), which had been sterilized with 70% ethanol and subsequently flushed with artificial cerebrospinal fluid, was then inserted through the craniotomy at a depth of either 2.6mm or 3.4mm to target the AON or PC. The insertion was performed with a manual hydraulic actuator (Narishige International). Once the catheter was lowered to the AON or PC, a first layer of 3-component dental cement (Super Bond C&B) was applied to secure the catheter to the skull and to cover the remaining space of the burr hole. Once the cement had fully cured (∼10 minutes) the catheter was firmly secured to the skull, so it could be released from the stereotactic frame. A set of fluidic fittings (compression sleeve – PostNova, CapTite One-Piece Fitting - LabSmith, CapTite Interconnect I - LabSmith) was mounted onto the tubing to secure a strong leak-free fluidic connection. More dental cement was used to secure these fittings to the skull and to cover the entirety of the incision and to make sure that there was no entry point for bacteria. Once all the previously applied cement was cured (∼10 minutes) the fluidic port was temporarily sealed with a screw-on cap (CapTite one-piece plug - LabSmith). The absence of air bubbles within the fluidic port was ensured by flooding the adapters with artificial cerebrospinal fluid (aCSF) prior to the fitting and sealing.

### **iv.)** Virus injection

For chemogenetic experiments, animals first received craniotomies bilaterally over the OB (AP 4.5 and 5, ML 0.75, DV 1.5). In each OB we injected 0.3µl of pENN.AAV.hSyn.HI.eGFP-Cre.WPRE.SV40 (retroAAV, Addgene). Two weeks later, a second surgery was performed to bilaterally target either the AON (AP 2.7, ML 1.35, DV 2.6) or aPCx (two injections per hemisphere: AP 2.34, ML 1.8, DV 2.75; AP 2.0, ML 2.2, DV 3.0). We injected at each cortical injection target, 0.3µl of pAAV-hSyn-DIO-hM4D(Gi)-mCherry (Addgene).

At the end of each surgical procedure, Atipamezole (Antisedan, 0.5mg/kg) was administered (IP), immediately followed by a dose of Meloxicam as analgesic (Metacam, 5-10 mg/kg) (IP). The animals were monitored during 5 days of recovery before performing any experiments.

### Odor delivery and behavioral monitoring

Odors were delivered with a custom-made, constant air flow olfactometer (Esquivelzeta Rabell et al., 2017). In short, an airflow passed through glass vials containing either mineral oil alone (non-odorized air, default state) or an odorant chemical dissolved in mineral oil (odorized air, odor stimulation). Each odorant was dissolved in paraffin oil to achieve a concentration of 1Pa partial vapor pressure. The odor filled flow was then delivered to the animal, straight in front of the nose and slightly angled downward to avoid airflow to the eyes. The air flow was kept constant during training and switched between mineral oil and odor conditions. Odor stimulation was 2s per trial.

Odors used were 2,3,5-trimethylpyrazine, 4-pentenoic acid, Acetic acid, Alpha-phellandrene, Anisole, Benzaldehyde, Benzyl acetate, Cinnamaldehyde, D-limonene, Ethyldecanoate, Ethyl valerate, Eugenol, Geraniol, Heptaldehyde, Heptanal, Iso-amyl acetate, Linalyl formate, Menthol, Methyl methacrylate, Octanoic acid, Phenylethyl alcohol, S+ carvone, Salicylaldehyde, Thiophene, Trans-2,cis-6-nonadienal (Sigma Aldrich). Odors were randomly assigned to novel or familiar categories and balanced between different animals to avoid possible biases from individual chemicals.

Respiration was recorded using an infrared camera (FLIR A325sc, 50mm lens, 60 fps, 320×240 pixels, uncooled, microbolometer), and inhalations were extracted using custom MATLAB scripts, as described earlier (Esquivelzeta Rabell et al., 2017; Mutlu et al., 2018).

### Behavioral paradigm

After a resting period of at least five days post-surgery, animals were habituated to handling and head-restrain during a period of 3-4 days. Mice were head-restrained in a light-proof, sound-attenuated metal box of 1m^3^. A red light was shining from above and odors were presented in front of the nose. During 4 subsequent training days (Fig. 2a), mice underwent one training session every day. In a single training session, mice were familiarized with 4 odorants, each presented at least 8 times (max 10 times), in random order, with an inter-trial interval (ITI) drawn from an exponential distribution with mean of 20s. The minimum and maximum ITI were constrained to 2s and 30s, respectively. After 4 days of familiarization, novelty sessions were performed. In the beginning of each novelty session, mice were randomly presented with 4 presentations of each familiar odorant used during the familiarization training. This pre-block was followed by the rest of the session with pseudorandomized delivery of 8 novel odorants, the 4 familiar odorants, and 2 blank conditions (mineral oil). Each odorant was presented at least 8 times (maximum 10 times). This experiment typically lasted about 1 hour 20 min. Since pharmacological agents become rapidly metabolized after injection into the brain, we limited the duration of the novelty session to ∼45 minutes in experiments involving convection enhanced delivery of muscimol or aCSF by presenting odors only 7 times.

### Electrode placement during acute electrophysiological recordings

To perform acute extracellular electrophysiological recordings, a mouse was head-restrained in the set-up. The silicone cap was cleaned with 70% ethanol and removed. The craniotomy was opened, the artificial dura was removed, and it was cleaned with sterile saline buffer. Neuronexus Poly 2 probes were inserted perpendicular to the surface of the exposed cortex. The highest channel located at white matter was employed as a reference. No artifacts were detected by the reference electrode from the spiking units. For acute recordings performed with Neuropixels 1.0 probes, the craniotomy opening was followed by the opening of the durotomy as well. When cleaned, a physiological saline solution (Physio-Sterop, Sterop) was put on top of the craniotomy. The probe was dyed with DiI (CellTracker CM-DiI, ThermoFisher Scientific) and inserted using a motorized micromanipulator (PatchStar, Scientifica). Insertion depth was determined using the Allen Brain Atlas (Allen Institute for Brain Science). After insertions of Neuropixels 1.0 probes, an additional layer of mineral oil was applied on top of the water reservoir. The probe was left for 20 to 30 minutes, the physiological solution on top of the craniotomy was replenished, and the recording and odor delivery were started. Recording sessions were initiated 20–30 min after probe placement and lasted 1–1.5 h. Probes were cleaned with 1% enzymatic solution (Tergazyme) for 30 min at the end of each recording session.

### Muscimol infusions

Chronically implanted microfabricated brain catheters were used for convection-enhanced delivery of muscimol (Muscimol BODIPY TMR-X, red fluorescence) or aCSF to the AON or PC. First, animals were placed in a head-restraining apparatus to allow for reliable access to the fluidic ports. The screw-on caps were removed and an infusion line leading to a syringe pump (Harvard Apparatus 33dds) was connected in its place. A low-volumes bolus of infusate was delivered at minimally invasive flow rates (0.025µl/min) whilst the inline pressure was monitored via a microfluidic pressure sensor (uProcess uPS0250-C360-10 – LabSmith). This monitoring allows the experiment’s interruption in case the pressure exceeds a certain threshold value to guarantee the animal’s safeguard. Upon the infusion’s termination, a 10-minute interval was used to allow for the rebalancing of pressure within the infusion system. After this interval the fitting was removed, and the screw-on plug was replaced. 30 min post-infusion, the behavioral experiment was started.

### Chemogenetic inactivation

To inactivate hM4D(Gi)-expressing neurons, the chemogenetic ligand clozapine-N-oxide (CNO; HelloBio, HB6149) was administered intraperitoneally at a dose of 2mg/kg (11.7 μl), diluted in saline. Injections were performed 30 minutes prior to the start of the behavioral paradigm. Control animals received an injection with an equivalent volume of saline. Animals were handled daily for at least five consecutive days before novelty experiments to habituate them to the injection procedure and minimize stress during experiments.

### Electrophysiological data acquisition

We used a DigiLynx (Neurolynx), Open Ephys and neuropixels recording systems (PXIe) to acquire electrophysiological signals at a sampling rate of 30 or 32 kHz respectively. Recordings were done using the ground screw as reference. For acquisition, we used Cheetah, Open Ephys and SpikeGLX softwares respectively.

### Histology

After behavioral experiments ended, animals were deeply anesthetized with ketamine-medetomidine, followed by transcranial perfusion with 0.1M phosphate-buffered saline (PBS) solution and 4% paraformaldehyde (PFA) solution. The brains were extracted and placed in PFA for 24 hours post-fixation. After post-fixation, brains were stored in PBS for 2 days. After those 2 days, the brain was sectioned in 60µm thick coronal slices with a vibratome (VT1000S, Leica). Free-floating sections were collected into 48 well plates filled with PBS. Sections from virus-injected animals were further processed for immunohistochemical staining. Coronal sections were incubated with primary antibodies chicken anti green (Molecular Probes) and rabbit anti red fluorescent proteins (Rockland) in PBT with 10% newborn calf serum (1:100 overnight 4°C); followed by two PBS washes and the subsequent incubation with secondary antibodies goat anti-chicken Alexa Fluor 488 and donkey anti-rabbit Alexa Fluor 568 (1:500 overnight 4°C, Molecular Probes) in PBT with 10% newborn calf serum. 3 PBS; we washed two times with PBS. All brain slices were mounted on glass microscopy slides with DAPI-containing mounting medium (4’,6-Diamidino-2-phenylindole dihydrochloride, Vectashield-H-1200-10, Sigma-Aldrich). Afterward, all brain slices were imaged by a confocal microscope (LSM 710 Zeiss).

### Calculation of co-expression efficiency

To assess the expression efficiency of the Cre-dependent virus (DREADDs-mCherry construct) with the Cre-expressing retro virus (expressing Cre), cells were manually counted across five different brain slices. Slices were randomly selected from either experiment targeting AON or aPCx cells projecting to the OB. For each slice, 50 cells were counted and classified as either fluorescently labeled with both GFP and mCherry signals or with mCherry signal alone. Co-expression efficiency (%) was calculated as the proportion of cells expressing both GFP and mCherry signals (GFP^+^ and mCherry^+^) relative to the total number of mCherry-positive cells (mCherry^+^), multiplied by 100.

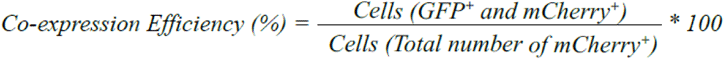

Cells expressing mCherry^+^ only indicated either leaky expression or the inability to detect GFP by immunohistochemistry.

### Estimation of total cell counts for AON and aPCx

To estimate the total number of labelled cells (mCherry, DREADDs-construct) in both the AON and aPCx, we used the following formula:

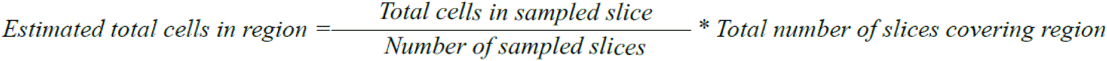

For each region, the number of cells in sampled slices was counted manually. 4 Different slices for the AON-OB projection dataset and 4 different slices for the aPCx-OB projection dataset, each from 4 different animals, were analyzed. Each region contained a total of 6 slices. The estimated total number of cells for each region was calculated by extrapolating the average number of cells per slice to the total number of slices covering the region.

### Extraction of inhalation onsets, calculation of baseline breathing frequency, and normalized breathing rate

The first inhalation onset after the odor onset was established as the stimulation onset and defined as time 0 s. Per trial, the breathing frequency was computed for each mouse in time bins of 50 ms. The binned average breathing frequency was computed per time bin per trial, whereby all inhalations in each time bin were combined and then divided by 50 ms. All values of each trial were then averaged to plot the breathing frequency (number of sniffs per second) against the different time bins. We normalized sniff data by subtracting the mean baseline before odor stimulation from each breathing frequency of the corresponding condition.

### Spike sorting and waveform characteristics and cell-odor pair selection

All recorded data were sorted using kilosort (https://github.com/MouseLand/Kilosort2). Individual units were then manually curated using phy (https://github.com/cortex-lab/phy). To ensure high-quality unit isolation, we only included neurons with a 0.9 presence ratio. The presence ratio was calculated by the number of seconds during which the unit was active (at least one spike) relative to the total length of the recording. In addition, we only included active neurons that had at least 3 spikes/s peak firing rate throughout the recording session. A drift contamination metric was calculated for each unit to evaluate the potential for contamination from non-stationarity over the data acquisition period. This metric estimates the sensitivity of the mean firing rate to random subsets of trials, and we included the units with 75% drift contamination. Cell-odor pair responsivity was visualized using a response index (2*auROC-1), comparing the distribution of odor-evoked spike counts to the pre-odor baseline. Cell-odor pairs were labeled as significantly odor-responsive using a rank-sum test (*p<*0.05) on trial-by-trial spike counts over the first sniff after odor delivery compared to spike counts over the last pre-odor sniff (Bolding et al., 2020).

### Modulation index

To quantify short term habituation invariant to the selectivity of a neuron, the mean firing rates for the first three and the last three trials of each odor presentation are represented as the x and y coordinates of a two-dimensional point cloud. To estimate the change relative to the unity line, a single-degree polynomial function is fitted to this point cloud. This best-fit line is constrained to pass through the origin (0,0). The slope of the constrained fit is then subtracted by 1 and multiplied by 100 to convert the values from ratios to percentages (Aydın et al., 2018).

### Neuronal distance to stable and multivariate linear regression

To assess changes in population activity during short-term habituation, we computed population firing rate vectors for each trial. These vectors represented the spiking activity of recorded neurons within two-time windows: the first was the 400 milliseconds following the first inhalation after odor presentation (odor trial), and the second was a control period spanning from 2400 to 2000 milliseconds prior to that first inhalation. To quantify the similarity of each trial’s population response to a stable baseline, we calculated the Euclidean distance between the population response of each trial and the mean response of the last three trials (stable trials). This distance was then normalized to the mean Euclidean distance between the stable trials themselves to account for inherent variability in the baseline. We therefore used the population distance-to-stable metric to quantify convergence of neuronal population responses toward a consistent state during short-term habituation (Bolding et al., 2020).

### Support Vector Machine SV

We used a linear Support Vector Machine (Kramer, 2016) for binary classification of novel and familiar stimuli from the firing rate of neurons recorded in AON or aPCx. Neurons from all sessions for each brain region were concatenated. To address the issue of overfitting, we used Leave-1-out cross-validation. Since sample sizes differed between regions, each iteration of the leave-one-out approach was averaged across different pools of decoded subpopulations. To allow for comparing the accuracy between regions, we subsampled the same number of neurons for each region (n=237). Over the course of 100 iterations, neurons were randomly selected from the entire pool. These neurons were then divided into separate sets for training and testing the SVM in each iteration. Finally, we balanced the novel and familiar trials in our dataset for the SVM since there were more novel than familiar stimuli in our experimental paradigm.

### Ordinary least squares linear regression

Ordinary least squares linear regression was used to quantify how much responses of each neuron were influenced by experimental variables. The full model included the following predictors: odor novelty, presentation number, interaction between odor novelty and presentation number, and the sniffing rate (in the window 0-500ms after the first inhalation). The dependent variable was the mean firing rate in each trial (0-400ms after the first inhalation). Because the predictors were not independent from each other (in particular, sniffing was higher for novel than for familiar odors), beta values could not be used to reliably estimate the contributions of individual variables. Instead, a series of reduced models - each excluding one or more predictors - was fitted and compared to the full model using F test. The p values obtained for all neurons were corrected for multiple comparisons using the FDR method (Benjamini and Hochberg, 1995). All analyses were performed using the statsmodels Python package.

### Tracking neurons across days

We extracted (chain) units that were stable over all days for each mouse via an Earth Mover Distance (EMD) based algorithm, proposed in (Yuan et al., 2024) using software from (https://github.com/janelia-TDHarrisLab/Yuan-Neuron_Tracking). The size of the resulting data was (number of chain units are in brackets): **AON**: MB107 (**49**), MB109 (**58**), MB118 (**7**), RDP009 (**2**); and aPCx: MB110 (**15**), MB111 (**54**), MB112 (**32**).

### Representational drift

To view the change in odor representations over days, we first computed odor representational similarity matrices, 𝑆𝑆_(𝑚𝑚, 𝑑𝑑), for each mouse, indexed 𝑚𝑚, on day 𝑑𝑑 . 𝑆𝑆_𝑚𝑚 = 𝑂𝑂_(𝑚𝑚, 𝑑𝑑) 〖𝑂𝑂_(𝑚𝑚, 𝑑𝑑)〗 ^𝑇𝑇, where 𝑂𝑂_(𝑚𝑚, 𝑑𝑑) is a matrix containing the representations of the odor set for mouse 𝑚𝑚 in rows on day 𝑑𝑑 and is of dimensionality 𝑛𝑛_(𝑜𝑜𝑑𝑑𝑜𝑜𝑜𝑜) × 𝑛𝑛_𝑢𝑢𝑛𝑛𝑢𝑢𝑢𝑢𝑢𝑢, where 𝑛𝑛_𝑜𝑜𝑑𝑑𝑜𝑜𝑜𝑜 = 4 is the fixed size of the odor set and 𝑛𝑛_𝑢𝑢𝑛𝑛𝑢𝑢𝑢𝑢𝑢𝑢 is the number of chain units for mouse 𝑚𝑚. To compare odor representational similarity matrices across days, we used the Centered Kernel Alignment (Kornblith et al., 2019) (CKA, with a linear kernel) between two representations, defined as:

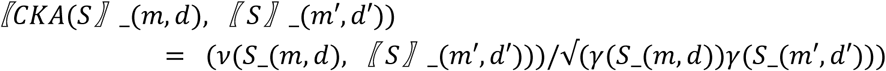

Where 𝜈𝜈(𝑆𝑆_(𝑚𝑚, 𝑑𝑑), 〖 𝑆𝑆〗_(𝑚𝑚′, 𝑑𝑑′)) is the effective number of shared dimensions between 𝑆𝑆_(𝑚𝑚, 𝑑𝑑) and

𝑆𝑆_(𝑚𝑚′, 𝑑𝑑′). 𝛾𝛾(𝑆𝑆_(𝑚𝑚, 𝑑𝑑)) is the effective number of dimensions (participation ratio) of the odorant representations 𝑆𝑆_(𝑚𝑚, 𝑑𝑑) [Giaffar 2024]. CKA is the ratio between the effective dimensionality of the (sub)space shared between datasets and the geometric mean of the individual effective dimensionalities of and is widely used as a measure of representational similarity in deep learning. A score of 1 indicates perfect alignment between representations (for a pair of days); a lower score reflects a decrease in the alignment, while accounting for differences in the effective dimensionality of neural responses over days. The CKA measures changes in the geometry of odor representations across days and can therefore be used to quantify this type of representational drift. Based on the CKA measure, we calculated the complementary CKA dissimilarity using the formula *CKA_DISS = 100*(1-CKA)*. MATLAB code to recreate these analyses are found in https://github.com/hamzagiaffar/090_AON_PCX_SIR.git

## Acknowledgements

This work was supported by Interne Fondsen KU Leuven C14/21/111 (SH), Fonds Wetenschappelijk Onderzoek-Vlaanderen (FWO-Flanders) fellowships 1135122N (ES), 1276122N (KK) and 1272222N (CA), and grant G097022N (SH).

## Author Contributions

Conceptualization and ideas: SH and CA; Experiments: RDP, MB; Analysis: HG, ES, KK, CA, RDP; Writing - Original Draft: SH; Writing - Review & Editing: RDP, MB, HG, ES, KK, CA, SH

## Data availability

Data will be made available upon request.

## Code availability

Code will be made available upon request.

## Correspondence and request for materials

Should be addressed to CA or SH

## Supplementary information

**Supplementary Fig. 1.**
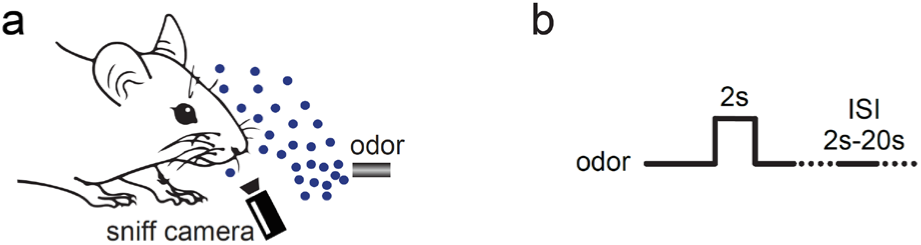
**a** Illustration of experimental setup. Odor delivery and an IR camera were directed towards the nose of the head-restrained animal **b** Odors were presented for 2 seconds with an inter-trial interval (ITI) drawn from an exponential distribution with mean of 20s. The minimum and maximum ITI were constrained to 2s and 30s, respectively

**Supplementary Fig. 2.**
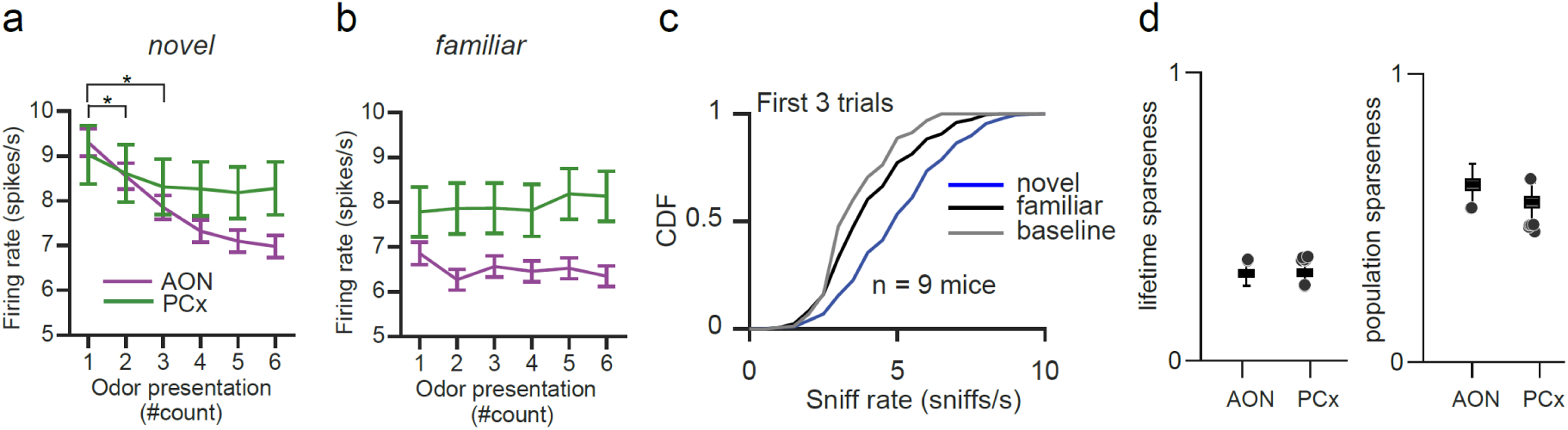
**a** Cumulative distribution comparison of mean sniff rates of first three presentations of novel (blue) and familiar (black) odors as well as baseline prior to odor stimulation (black) activity. **b** Population habituation to novel odors in AON (purple) and aPCx (green). **b** Sames as (a) for familiar odors. **d** Comparable lifetime (left) and population (right) sparseness between AON and aPCx neurons, each dot represents outliers of the mean distribution to novel odors.

**Supplementary Fig. 3.**
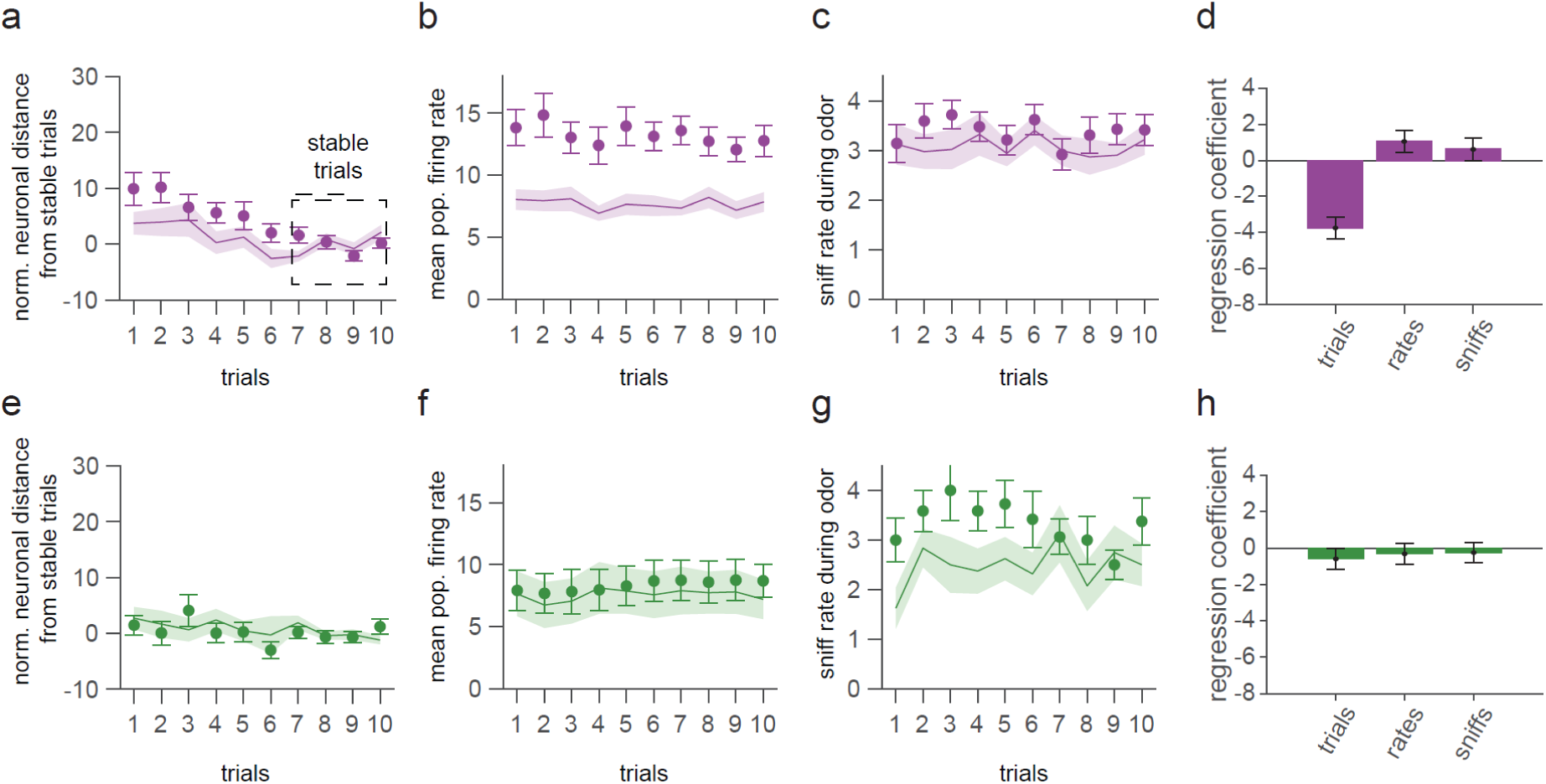
**a** Average Euclidean distance between familiar trial population vectors and stable trials in AON (shading indicates ± SEM). Baseline activity prior to odor presentation is represented as continuous line. **b** Average population firing rates to familiar odors in AON. Baseline activity prior to odor presentation is represented as continuous line. **c** Average AON sniff rates to familiar odors (shading indicates ± SEM). Baseline respiration rates prior to odor presentation are represented as continuous line. **d** Multiple linear regression coefficients for effects of trial number, population firing rate, sniff rate (all not significant) on population distance to stable in AON (mean ± SEM). **e-g** Same as (a-c) for aPCx. **h** Multiple linear regression coefficients for effects of trial number, population firing rate, sniff rate (all not significant) on population distance to stable in aPCx (mean ± SEM).

**Supplementary Fig. 5.**
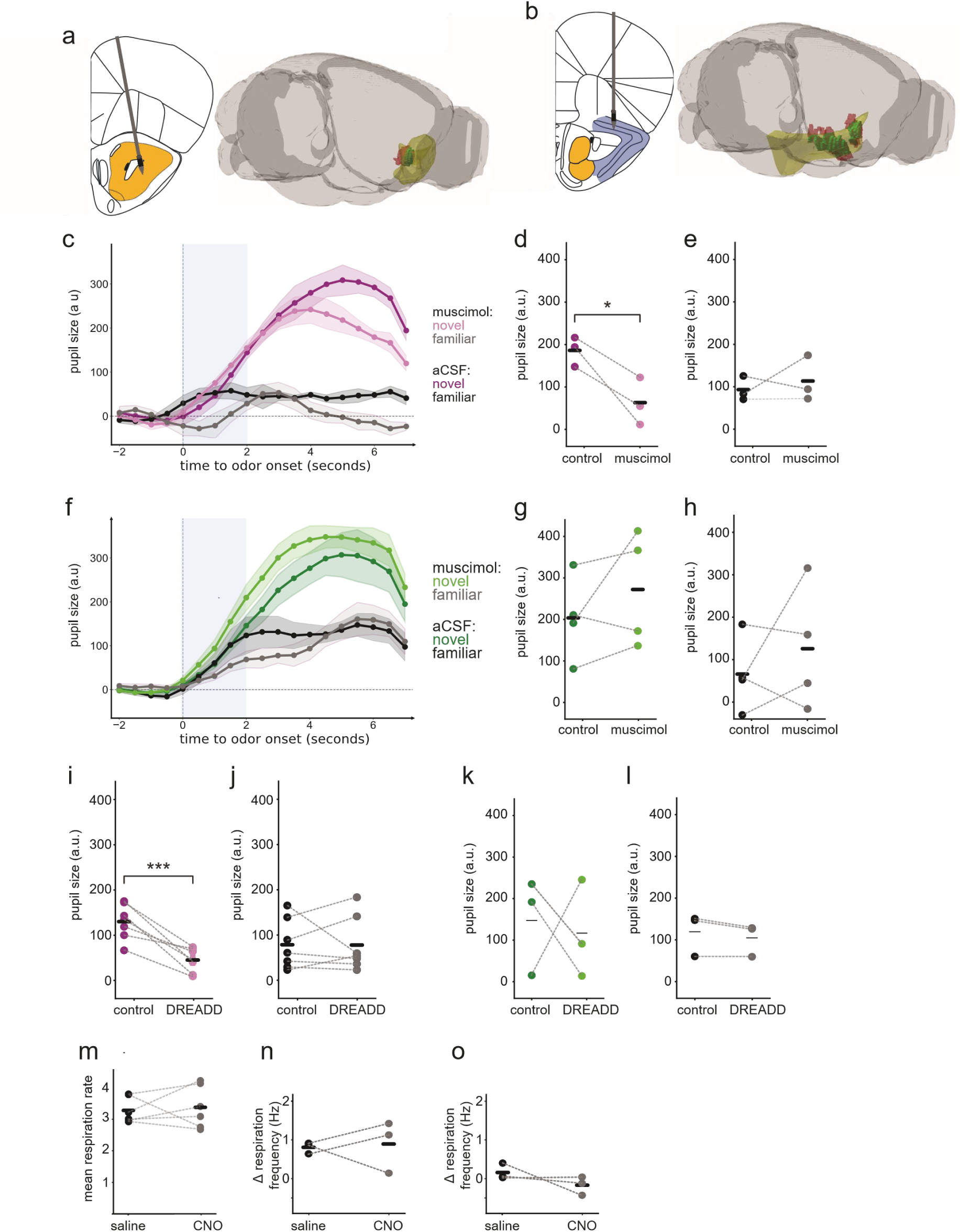
**a** (**left)** Implanted microfabricated brain catheter in the target region AON **(right)** Example reconstruction of the delivery cloud after infusion performed in the AON (yellow) with on-target volume (green, ∼37% of AON is targeted) and off-target volume (red). **b (left)** Implanted microfabricated brain catheter in the target region, aPCx. **(right)** Reconstruction of the delivery cloud for the infusion performed in the aPCx (yellow) with on-target volume (green, ∼40% of the anterior section of the aPCx is targeted) and off-target volume (red). **c** Normalized change in pupil size during the first three trials (mean ± SEM) after AON injection with muscimol (novel trials: light purple, familiar trials: light grey) or aCSF (novel trials: dark purple, familiar trials: dark grey, n=3 mice). Blue shading represents the odor stimulation window. **d** Mean pupillary response to novel stimuli after injection of muscimol or aCSF in the AON (paired t-test, \**p*=0.04). **e** Same as (b) for first three familiar trials (not significant). **f** Normalized change in pupil size during the first three trials (mean ± SEM) after aPCx injection with muscimol (novel trials: light green, familiar trials: light grey) or aCSF (novel trials: dark green, familiar trials: dark grey, n=4 mice). Blue shading represents the odor stimulation window. **g** Mean pupillary response to novel stimuli after injection of muscimol or aCSF in the aPCx (not significant). **h** Same as (g) for first three familiar trials (not significant). **i** Mean pupillary response to novel stimuli after injection of CNO or saline in animals with DREADD virus targeted to AON (n=7 mice; paired t-test, \*\*\**p*=0.001). **j** Same as (i) for first three familiar trials (not significant). **k** Mean pupillary response to novel stimuli after injection of CNO or saline in animals with DREADD virus targeted to aPCx (n=3 mice; not significant). **l** Same as (k) for first three familiar trials (not significant). **m** Mean baseline respiration rate in mice not expressing the DREADD receptor but injected with saline or CNO (n=6 mice; not significant). **n** Mean sniffing response to novel stimuli after injection of saline or CNO (n=3 mice; not significant). **o** Same as (n) for first three familiar trials (not significant).

